# Modeling The Effect of Background Sounds on Human Focus Using Brain Decoding Technology

**DOI:** 10.1101/2021.04.02.438269

**Authors:** Aia Haruvi, Ronen Kopito, Noa Brande-Eilat, Shai Kalev, Eitan Kay, Daniel Furman

## Abstract

The goal of this study was to investigate the effect of sounds on human focus and to identify the properties that contribute most to increasing and decreasing focus in people within their natural, everyday environment. Participants (N=62, 18-65y) performed various tasks on a tablet computer while listening to either no background sounds (silence), popular music playlists designed to increase focus (pre-recorded songs in a particular sequence), or engineered soundscapes that were personalized to individual listeners (digital audio composed in real-time based on input parameters such as heart rate, time of day, location, etc.). Sounds were delivered to participants through headphones while simultaneously their brain signals were recorded by a portable electroencephalography headband. Participants completed four one-hour long sessions at home during which different sound content played continuously. Using brain decoding technology, we obtained individual participant focus levels over time and used this data to analyze the effects of various properties of sound. We found that while participants were working, personalized soundscapes increased their focus significantly above silence (p=0.008), while music playlists did not have a significant effect. For the young adult demographic (18-36y), all sound content tested was significantly better than silence at producing focus (p=0.001-0.009). Personalized soundscapes increased focus the most relative to silence, but playlists of pre-recorded songs also increased focus significantly during specific time intervals. Ultimately we found that it is possible to accurately predict human focus levels that will be experienced in response to sounds *a priori* based on the sound’s physical properties. We then applied this finding to compare between music genres and revealed that classical music, engineered soundscapes, and natural sounds were the best genres for increasing focus, while pop and hip-hop were the worst. These insights can enable human and artificial intelligence composers to produce increases or decreases in listener focus with high temporal (millisecond) precision. Future research will include real-time adaptation of sound libraries for other functional objectives beyond affecting focus, such as affecting listener enjoyment, stress, and memory.

## 1 Introduction

### 1.1 The effect of sound on human experience

Sounds are all around us, from natural sounds like the wind, to engineered sounds like music. It is well-established that sounds have a major influence on the human brain and consequently, human experience (Levitin, 2006; Sacks, 2010). Sounds can reduce stress (Davis & Thaut, 1989), support learning and memory formation (Hallam et al., 2002), improve mood (Chanda & Levitin, 2013), and increase motivation (Salimpoor et al., 2015). Sounds can also do the opposite and create aversive experiences (Kumar et al., 2012; Schreiber & Kahneman, 2000; Zald & Pardo, 2002). One of the most significant effects of sounds is to impact focus. Focus is commonly demanded by tasks of daily living and work, and in these areas sounds offer a safe way to increase focus levels and productivity. However, sounds can be beneficial or distracting, and previous results have been inconclusive in determining the reason (de la Mora Velasco & Hirumi, 2020).

For example, it has been found that listening to music with lyrics while reading or working can decrease concentration or cognitive performance (H. Liu et al., 2021; Shih et al., 2012), while several studies have shown oppositely that natural-occurring sounds such as white noise, as well as classical music, can be beneficial for increasing focus and can improve learning outcomes (Angwin et al., 2017; Chou, 2010; Davies, 2000; Gao et al., 2020). Therefore, one interesting question emerges which is: what are the specific properties of sounds that affect human focus levels the most? Additionally, studies have shown that the effect of sounds is often subjective, where whether one likes a given sound or not is a key factor in its effect on their experience (Cassidy & Macdonald, 2009; Huang & Shih, 2011; Mori et al., 2014). Although this finding about the subjectivity of sound reappears across many studies, psychophysical thresholds are known to exist and there are clearly natural laws governing much of the way humans hear and experience sound (Levitin et al., 2012; Nia et al., 2015; Washburne, 2020).

The potential of sounds to increase focus and demand for non-pharmaceutical tools that enable individuals to enhance their ability to focus has recently led several companies (including Endel, Brain.fm, Mubert, Enophone, Focus@Will, Melodia, AIVA, and others) to develop soundscapes that are dedicated to increasing focus on-demand. These soundscapes include elements of white noise, music, and other sonic properties that are functionally combined to increase a listener’s focus and maintain high levels of focus over long durations of time. One of the challenges in this field is to figure out the physical properties of sound that contribute to human experience the most so that design principles can be defined correctly to create soundscapes that achieve the goal of increasing focus, opposed to the inverse of causing distractions and hurting an individual’s ability to focus. Insights about sound properties therefore have been sought by commercial groups alongside academic groups in order to learn how to optimize experiences through sound.

Many scientific studies have explored this question and looked for the relationship between sound, music and human experience using objective measures that empirically assess properties of audio and their emotional correlates. For example, Cheung et al (Cheung et al., 2019) found that pleasure from music depends on states of expectation, such as a skipped rhythmic beat, which can either be pleasurable or discomforting depending on the listener’s circumstance. *Sweet Anticipation* (Huron, 2006) similarly maps how music evokes emotions within a theory of expectation and describes psychological mechanisms that are responsible for many people’s mixed responses to sounds. Other studies used machine learning methods to map from features of audio signals to emotions (Brotzer et al., 2019; Cunningham et al., 2020; Hizlisoy et al., 2021; Vempala & Russo, 2012; Yang et al., 2008). These machine learning studies to date have, however, only aimed to predict emotions based on the limited valence-arousal circumplex model, and as far as we know, no attempts have been made to predict human focus levels exclusively based on audio signal analysis.

One persistent obstacle to the field’s understanding has been studies that rely on data with a low temporal resolution. Since sounds and emotions evolve fast, on the order of tens of milliseconds, the current lack of modeling tools capable of capturing the fast changes in human experience that accompany changes in sound is a major hindrance to progress (Cowen & Keltner, 2017; Larsen & Diener, 1992). Commonly, for example, reports are based on data where there is a single emotional label per song, while the song lasts ~2-3 minutes and throughout it there are emotional dynamics that change dramatically. This mismatch of data can lead to conclusions being drawn from inadequately small amounts of samples, and worse than that, inaccurate emotional labels.

### 1.2 Attention and emotion decoding from brain signal

Brain decoding technology offers an exceptional opportunity to tackle this issue, since it enables us to get an estimation for the experience dynamics at the same time resolution as focus phenomena occur. Using electroencephalogram (EEG) sensor data, which contains electrical brain activity measured from the scalp (non-invasive) on the order of hundreds of measurements per second, many studies have established that it is possible to capture fast changes in human emotions and experience, such as stress (Perez-Valero et al., 2021), arousal (Faller et al., 2019), fatigue (Hu, 2017), and happiness (Lin et al., 2017). Several studies have similarly shown the ability to capture focus and attentional state changes, affirming that this information as well is captured in EEG sensor data (Hamadicharef et al., 2009, 2009; Jung et al., 1997; Micoulaud-Franchi et al., 2014; Tuckute et al., 2021). While brain decoding technology has been applied widely to study the effects of different types of stimuli (e.g visual, tactile, auditory) on human experience (Asif et al., 2019; Bhatti et al., 2016; Shahabi & Moghimi, 2016), as far as we know, it has not been applied to study the joint effects of sound and focus at the high temporal resolution needed to explain both phenomena.

In recent years, progress in the development of non-clinical, wearable EEG sensors (such as Muse, Neurosky, Emotiv, Bitbrain, etc.), which are intended for consumer uses, has led to new research paradigms where comfortable, affordable, wireless, and easy-to-use at-home measurement devices collect neuroscientific data “in-the-wild” at a large scale and make it possible for the first time to measure brain responses from diverse audiences within their natural, real-world environment. Many of the wearable devices offer decoding outputs beyond the raw sensor data, and these “off-the-shelf” decoding outputs include attention, relaxation, and other states (Abiri et al., 2019; Bird et al., 2019; González et al., 2015; N.-H. Liu et al., 2013; Rebolledo-Mendez et al., 2009). It is important to note, however, that although decoder algorithms exist in the market for consumer uses, verifying their reliability to accurately capture attention, valence, arousal, stress and other attributes of human experience at a high temporal resolution and research quality has remained a challenge.

### 1.3 Combining brain decoding with sound tests to increase focus

In the current study, we used Arctop’s brain decoding technology (neuOS^TM^) on data from portable EEG (Muse-S) headbands to measure human focus levels in individuals performing tasks at home while listening to different types of sounds. Since this is a relatively new decoding technology, we first evaluate the validity of the focus outputs within the experimental conditions. Then, once convinced of the output’s veracity and reliability, we use the focus data to compare effects of different sound stimuli on individuals while performing different tasks.

Next, we exploit the decoded data’s high time resolution to map between raw audio signals and the focus dynamics. Based on this mapping, we build a model that takes sound properties and predicts human focus levels, enabling us to compare between new songs, sounds, and between genres to gain additional insights about the nature of sounds which drives human focus the most. These insights can help in the future to generate optimal playlists to increase focus, engineer better soundscapes, and even adapt sounds in real-time based on an individual’s focus levels to enable them to precisely influence their own mental state.

## 2 Materials and Methods

### 2.1 Participants

Sixty-two (62) participants (40 males, 22 females, 18-65 years), completed four (4) sessions over a single (1) week at their own home. All participants were recruited from an opt-in screening panel and were distributed across the five (5) major regions of the continental United States (Northeast, Southwest, West, Southeast, and Midwest). Only participants who reported normal hearing, normal vision, or vision that was corrected to normal with contact lenses, were included. We excluded volunteers who reported using medication that might influence the experiment and who reported neurological or psychiatric conditions that could influence the results. Participants were native English speakers and a written informed consent was obtained from each participant prior to their participation. Participants received compensation for their time.

### 2.2 Paradigm

#### 2.2.1 Tasks

Participants performed various tasks within a mobile app (neuOS^TM^ by Arctop Inc.) while listening to one of three types of sound and wearing a brain signal measuring headband (4-channel EEG Muse-S device by Interaxon Inc.). Each participant received a kit at their home that included all the equipment needed to participate, including over-ear (Sony) headphones, headband and tablet computer with the mobile app installed. Participants recorded four one hour long sessions, while listening to different types of sounds. Sessions included 30 minutes of a “Preferred Task” — a task chosen by the participant — followed by short tasks used to validate the brain decoding outputs for each session. These validation tasks included video games (Tetris), math problems (Arithmetics), and word problems (Creativity). Participants were assigned to groups according to a pseudorandom schedule that controlled for potential sequence effects of the tasks and different sound stimulus types (Fig. 1). The short tasks were used to calibrate the Arctop decoding algorithms to a validated performance level, and afterwards the validated model was used to measure each participant’s focus level across the Preferred Task.

**Figure 1.**
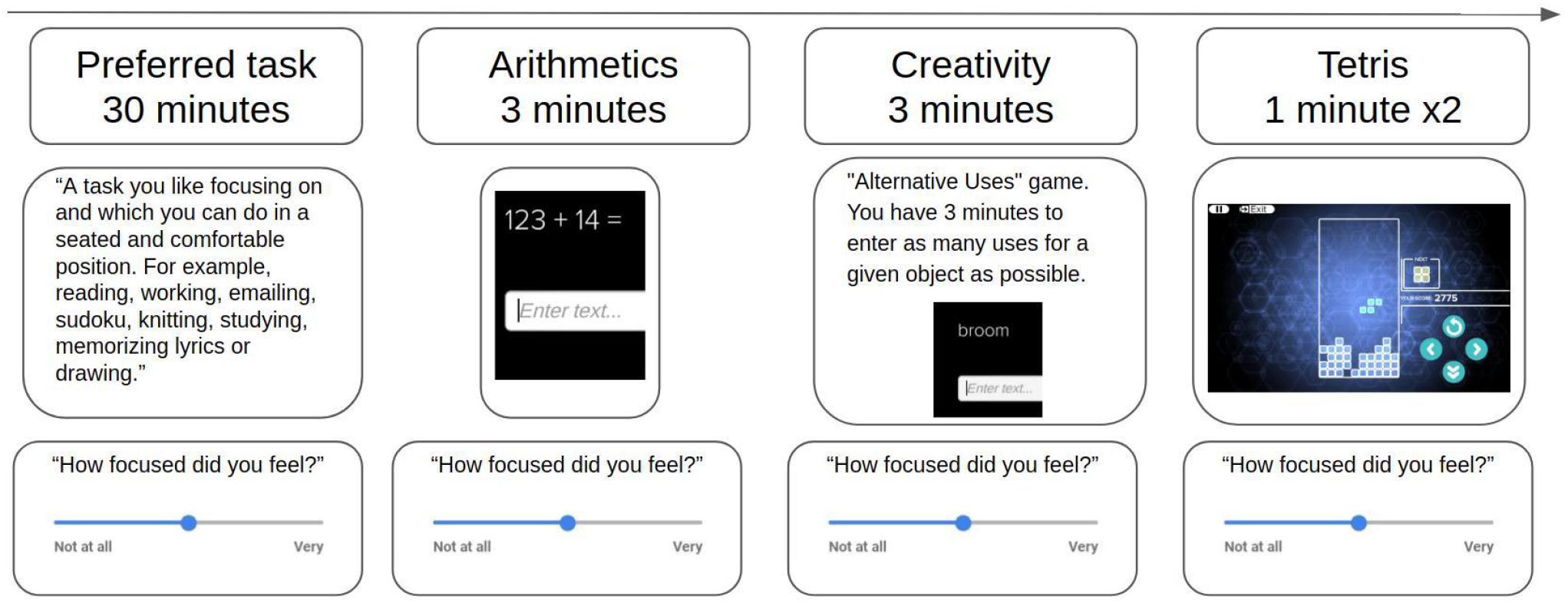
Schematic illustration of the paradigm in each recording session. Each session started with 30 minutes of a task selected by the participant (“Preferred Task”), followed by 3 minutes of arithmetics exercises, 3 minutes of a creativity task, and two levels of Tetris the video game (each level lasted 1 minute regardless of performance). After each task, participants answered a survey where they reported on aspects of their experience (e.g. focus, enjoyment, stress) using linear scale sliders from “Not at all” (0) to “Very” (1).

Participants were instructed to choose a Preferred Task they could perform in a seated position while listening to sounds through the headphones, and which they would be happy to repeat in all four sessions. For example, Preferred Tasks that were chosen included knitting, working, reading, and solving Sudoku puzzles. At the end of each task the participants self-reported their experience through a survey in the app which used linearly-scaled slider buttons to quantify experience along several dimensions (e.g. focus level, enjoyment, stress, motivation, etc.). For the Preferred Task, the survey included reporting on their focus level during the first and second half of the task separately, resulting in six (6) self-reported quantitative focus labels per session (Preferred Task: 2 labels, arithmetics: 1 label, creativity: 1 label, tetris: 2 labels).

#### 2.2.2 Sounds

Each participant experienced four sound conditions over the four days of the study: two music playlists by leading digital service providers Spotify and Apple (downloaded September 2020), one personalized soundscape engineered by Endel, and silence (no audible sounds). We selected Spotify’s ‘Focus Flow’ playlist and Apple Music’s ‘Pure Focus’ playlist to represent the category of pre-recorded sounds designed to increase focus. For soundscapes we selected the mobile application Endel to represent the category of real-time, engineered sounds that contain a mixture of noise and musical properties. The Endel app ‘Focus’ soundscape was used by each participant on their own device. All sound conditions were instrumental (i.e. did not include singing or any audible lyrics). For the condition of silence, participants wore headphones exactly as they did in the sound conditions, but no music or audible sounds of any kind were played and no soundscape was generated - participants simply completed the session in a quiet environment.

### 2.3 Data processing

#### 2.3.1 Data acquisition

While participants were listening to sounds and engaging in the experimental tasks, their electrical brain activity was recorded using a portable, noninvasive electroencephalograph (EEG) headband that weighed 41 grams (Muse-S device by Interaxon Inc). The headband included four dry fabric EEG sensors (sampling rate: 256 Hz), photoplethysmography (PPG) sensors (for heart rate) and motion sensors (gyroscope, accelerometer). The brain-measuring EEG sensors are located on the scalp at two frontal channels (AF7, AF8) and two temporal channels (TP9, TP10), with the reference channel at Fpz. The headbands were put on by participants themselves with the assistance of a quality control screen that started each session by giving participants real-time feedback on the signal quality and made it easy to adjust the headband appropriately to acquire an optimal signal quality (Fig. 2). No technicians or other support staff assisted in the placement of the headbands - the process was completely automated by the in-app prompts, freeing the participants to complete sessions at any time of their choosing.

**Figure 2.**
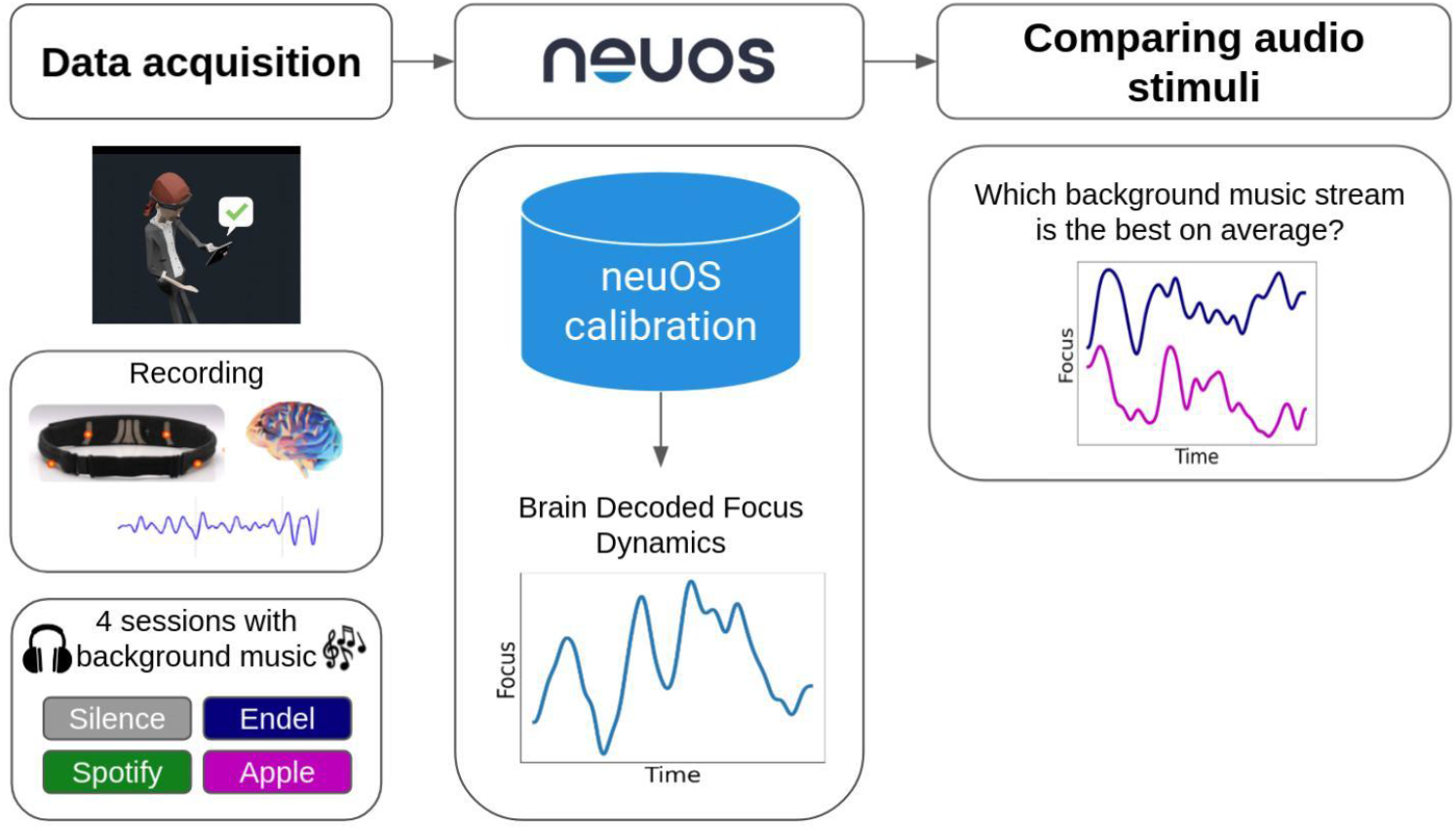
Schematic illustration of the data processing pipeline. Data acquisition included at-home recordings of four sessions, each with a different background sound type. Arctop’s neuOS brain decoding technology was used to predict the focus dynamics at a rate of 5Hz. Obtaining the brain decoded focus dynamics synchronously with the sound content enables comparison of focus levels correlated with different physical properties of sound.

#### 2.3.2 Brain data based models of focus

Arctop brain decoding technology (neuOS) was used to transform the sensor data into predicted focus dynamics with a time resolution of 5Hz (Fig 2). For each participant, short tasks (games, word and math problems) were used to calibrate and validate a model of their focus based on the brain data, and then once validated the model was applied to the Preferred Task data. Fig. 3 shows the resulting brain decoded focus levels of two representative participants across all four sessions during the Preferred Task. Model performance was evaluated using Pearson correlation coefficient between the self-reported focus and the brain decoded focus values, and after thresholding the values, with the area under the ROC curve for binary classification of low/high focus (Fig. 5). Eleven (11) participants were excluded from further analysis following model validation due to excessive noise in their recorded brain data and/or unreliable survey responses, leaving a total of 51 participants (mean age= 36, SD=8, 17 females and 34 males) in the experimental analysis.

**Figure 3.**
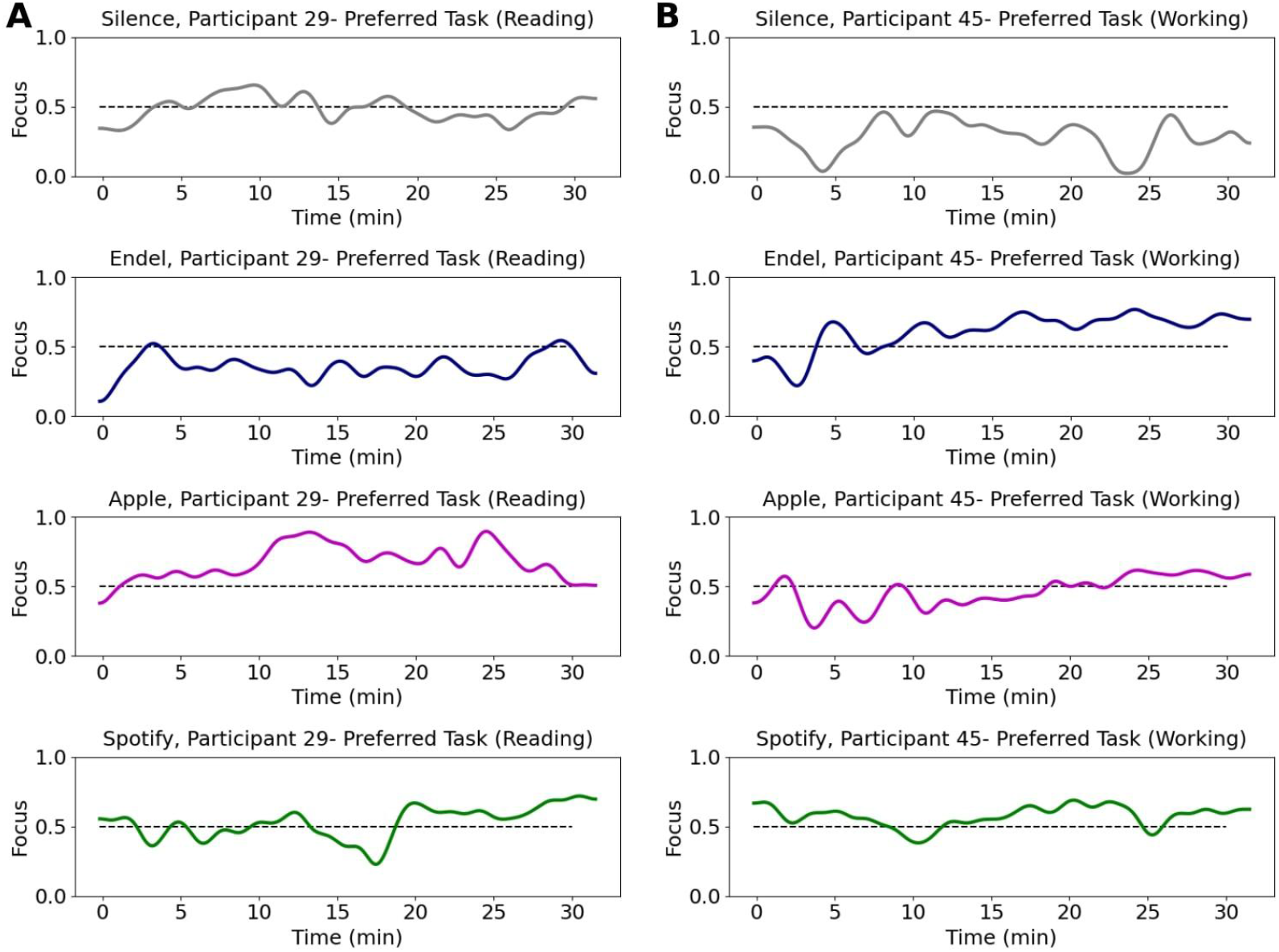
Brain data based focus model dynamics of two representative participants during the Preferred Task performed at each of the four sessions. Each row represents a session with a different sound stream playing in the background as participants perform their chosen task. Each session included 30 minutes (X axis = time in minutes) of a “Preferred Task” over which their focus level (Y axis = decoded focus) was measured. Participant 29 **(A)** was reading while Participant 45 **(B)** was working.

#### 2.3.3 Statistical methods

For comparisons between average focus levels during the different sound content presented, we calculated for each participant (N=51) the median focus level while performing the preferred task and conducted a one-way repeated measures ANOVA (Analysis of variance) test. Then, if p<0.05, paired t-tests were applied post hoc to compare between pairs of sound streams using Holm-Bonferroni correction. Time series statistical tests were applied to compare focus level dynamics and discover specific time periods of significant difference. A paired t-test was applied at each second between focus levels of two sound streams. The p-values were then corrected for multiple comparisons by setting a threshold for a minimum significant sequential time-samples. The threshold was determined by random permutations (1000 iterations) of participants’ conditions and repeating the statistical test, resulting in a distribution of significant sequential time samples. The threshold was set as the 95% percentile of the resultant distribution (Broday-Dvir et al., 2018).

#### 2.3.4 Sound signal decomposition and feature extraction

The pre-recorded playlists conditions (Apple and Spotify) provided raw sound data that we used to obtain sound property dynamics in the time and frequency domain that could be correlated with the obtained focus dynamics. Soundscapes were not used in this analysis because they were produced in real-time personally for each participant, which limited the ability to apply sound property analysis appropriately across the data set. The sound features were calculated using Python’s library pyAudioAnalysis (Giannakopoulos, 2015), for example, the sound signal energy, spectral entropy, and chroma coefficients. The features were calculated in short-time windows of 50 ms with a sliding window of 25 ms. Then, basic statistics were calculated over the sound features in windows of 30 seconds (e.g. mean and std), resulting in 136 sound properties ([link to full list](https://github.com/tyiannak/pyAudioAnalysis/wiki/3.-Feature-Extraction)). To enable mapping to the brain model, the brain decoded focus levels were also averaged in the corresponding 30 seconds windows (Fig. 4).

**Figure 4.**
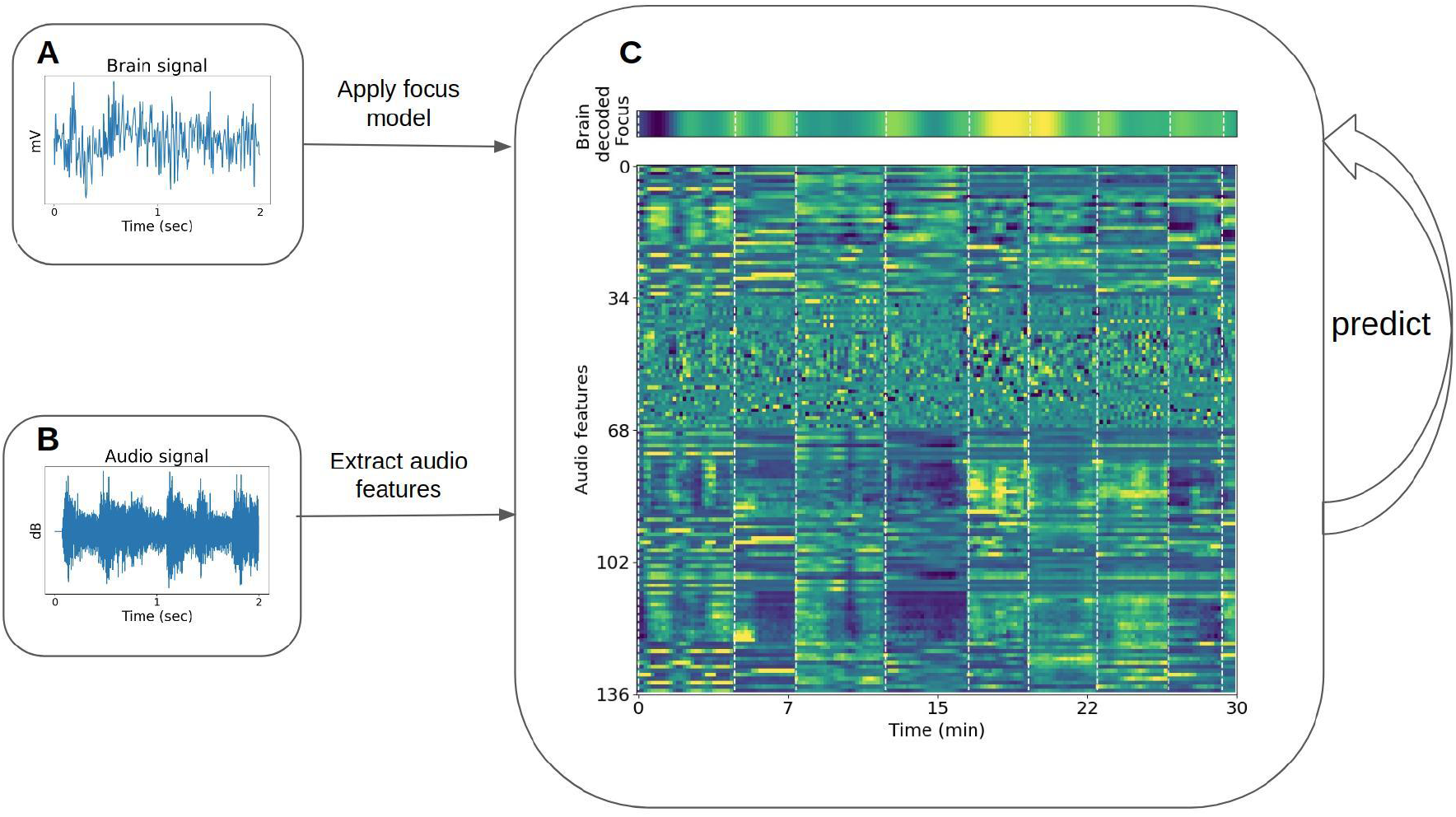
Diagram demonstrating the framework for correlation of time-series focus values with sound properties. **(A)** Example of a recorded brain data in microvolts (single channel of EEG) segment, which after applying the preprocessing and trained models on 30 minutes of recordings, transforms to the brain decoded focus dynamics (**top (C)**). **(B)** Examples of a sound segment in decibels taken from one of the songs. **Bottom (C)** The sound features (y-axis) dynamics during 30 minutes of recordings (x-axis).

To obtain the threshold for significant correlations between sound features and focus levels (p<0.05), a shuffle analysis was performed. Random permutations (1000 iterations) of the brain decoded focus levels were applied across songs to preserve the time dependency of focus levels within a song and the focus levels distribution. The correlation of each sound feature was calculated with the permuted focus level. The threshold was set as the 95% percentile of the resulting correlation’s distribution.

#### 2.3.5 Obtaining the sound decoded focus model

To map the relationship between properties of the sounds heard and focus levels measured from the brain, we first applied principal component analysis (PCA) to reduce the dimensionality of the sound features. We then trained regression models between the transformed sound features and the brain decoded focus through a 5-fold cross validation procedure that used 80% of the songs in each iteration to train and 20% to test. The presented sound decoded focus model is a linear model based on the first PCA component of the features (shifted and rescaled).

## 3 Results

### 3.1 Brain-measured focus levels accurately reflect self-reported focus levels

Before comparing focus levels elicited by the different sound types, we validated the underlying brain decoding technology by comparing between the brain-based focus predictions and the self-reported focus values. Figure 5A shows a histogram of the model performance per participant. The model is evaluated based on the AUC score (of the ROC curve) for prediction of self-reported focus during the Preferred Task (low-high focus) where the chance guessing level is = 0.5 (black dashed line). The average result across participants obtained was <AUC>=0.83 (N=51, SD=0.19), a strong validation of the brain-measured focus model’s accuracy.

**Figure 5.**
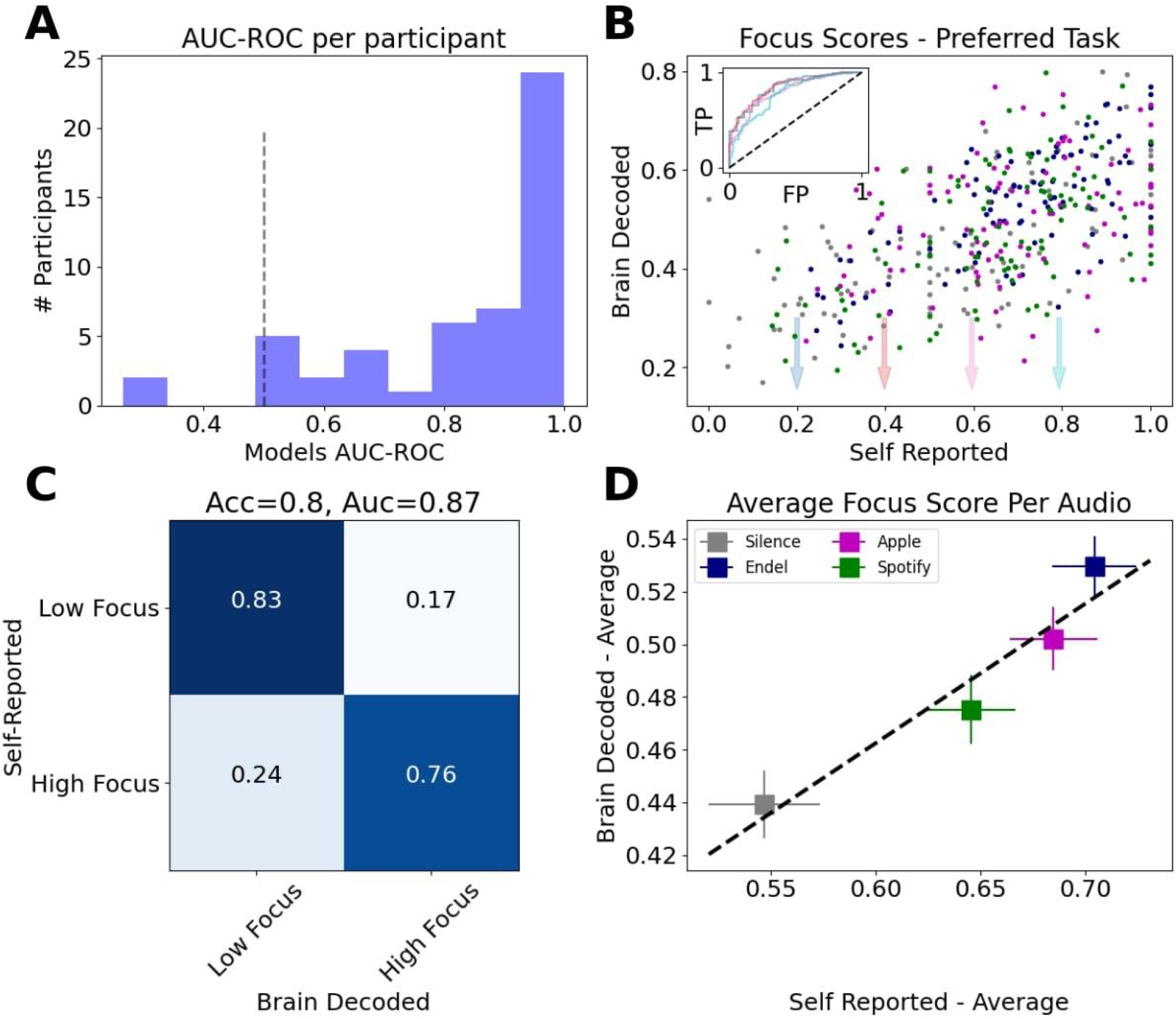
Validation of focus measurements derived from brain data. **(A)** Histogram of focus models performance per participant (N=51), evaluated using the area under the ROC curve (AUC-ROC). Black dashed line marks chance level (0.5). **(B)** Average focus levels per event vs. self-reported focus resulted in Pearson correlation of 0.6. Inset shows ROC curves for different values of self-report threshold. **(C)** Confusion matrix after thresholding the focus score predictions and self-report. Classification scores for 2-classes (low focus vs. high focus) are AUC=0.87 (area under ROC curve), Accuracy=0.8. **(D)** Average brain decoded focus levels vs. average self-reported focus across the four sound types.

When aggregating the tasks from all participants, the Pearson correlation between the brain decoded focus model and the self-reported focus was Corr(414)=0.6, p<10^−4^ (Fig. 5B). The inset in Figure 5B shows the ROC curves for different values of self-reported threshold and the confusion matrix for one of these thresholds (0.4) resulted in an accuracy score of 0.8 (Fig. 5C). Figure 5D shows the average brain decoded focus level per sound type vs. the average self-reported score.

### 3.2 Soundscapes induce a higher focus level compared to silence

Using the validated focus models which output five measurements per second (5Hz), we then compared between the average focus levels elicited by the sounds during the Preferred Task. The background sound was found to have a significant effect (top row in Table 1, F(3,150)=4.28, p=0.006, statistical methods for details) on the elicited focus level, and the post hoc tests (Holm-Bonferroni correction) revealed that streaming soundscapes (with Endel app) was significantly higher compared to silence (Fig. 6A1, supp. Table 1, M=0.090, SE=0.027, t(50)=-3.38, p=0.008), while streaming music using Apple or Spotify did not have an effect (Apple: t(50)=-2.37, p=0.11, Spotify: t(50)=-1.24, p=0.65). For 35.3% of the participants the Endel session produced their highest focus level, while for 27.5% of participants the Apple playlist produced the highest focus level. For 19.6% of participants Spotify was best for producing focus and for 17.6% silence was (Fig. 6A2, the details sorted focus levels per participant are shown in Supp. Fig. 1).

**Table 1.** Results of a one-way repeated measures ANOVA performed on each subgroup comparing the average brain decoded focus levels of each sound stream during the Preferred Task. Sound most significantly affected those below 36 years old.

| Group | N | F | p |
| --- | --- | --- | --- |
| All | 51 | 4.28 (3,150) | 0.006 |
| Working | 26 | 3.74 (3,75) | 0.014 |
| Not Working | 25 | 1.91 (3,72) | 0.14 |
| Age>36 | 26 | 1.81 (3,75) | 0.15 |
| Age<36 | 25 | 6.97 (3,72) | <0.001 |

**Figure 6.**
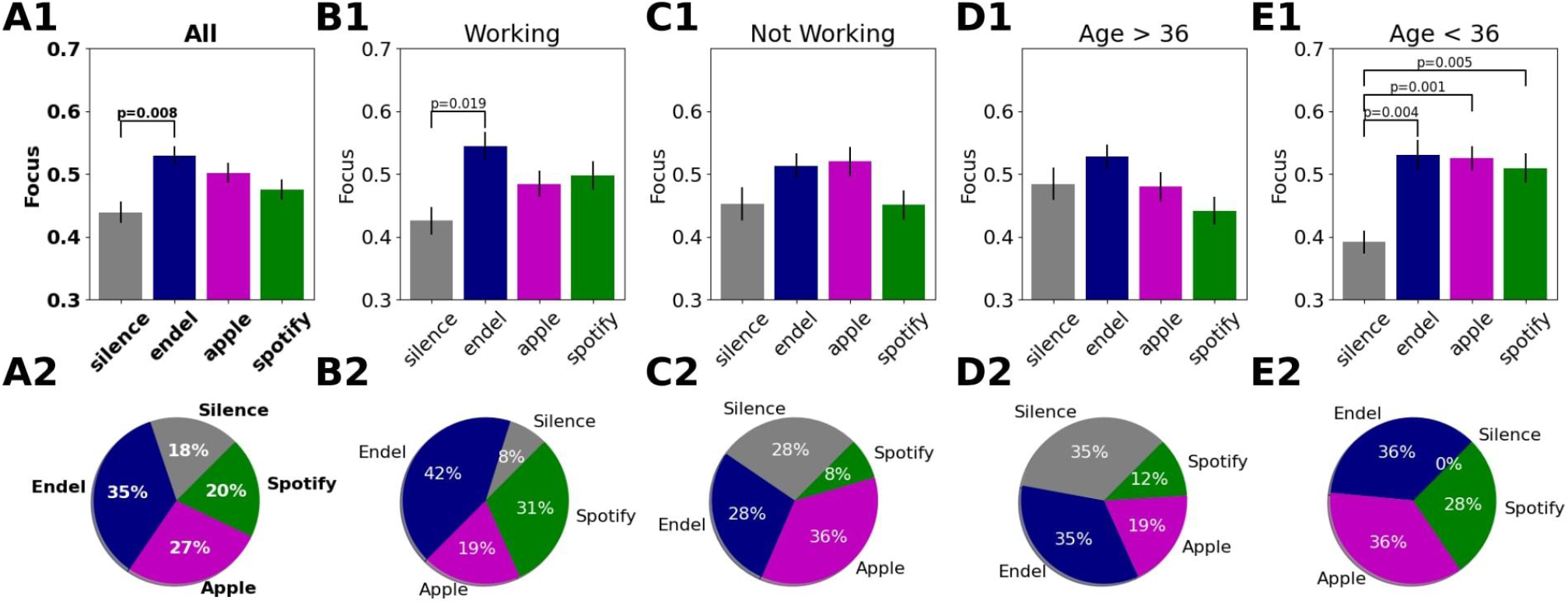
Comparison of the brain decoded focus during the Preferred Task while listening to different sounds. Top row - Average focus levels for each sound stream during the Preferred Task for each group of interest, including statistical results. Error bars are standard errors. Bottom row - Distribution of the best session (highest focus on average) for each participant per group. The groups of interest are: **(A)** All participants (51), **(B)** Participants who were working during the Preferred Task (26), **(C**) Participants who were not working (25 -reading, knitting, playing, etc). **(D)** Participants above 36 (26). **(E)** Participants below 36 (25).

To gain a better understanding of the conditions where sound affected focus, we next split the participants into subgroups of interest and repeated the statistical analysis. We first asked whether the focus level difference is task dependent. During the Preferred Task, 51% of the participants (26) chose to work, while the rest (49%) read a book (29.4%), played games (9.8%) or did other various tasks (e.g. knitting, 9.8%). To assess the effect of sounds on focus levels during these different tasks, we split the participants to the ones who worked and those that did other tasks. We found that for the “working” group, the focus level elicited by Endel’s soundscapes was higher compared to silence (M=0.12, SE=0.04, t(25)=3.26, p=0.017), while for the “not-working” group there was no difference (Fig. 6, supp. Table 1, M=0.06, SE=0.04, t(24)=1.552, p=0.447). These results suggest that the focus level differences between Endel and Silence are task-dependent, where the sound is beneficial for specific types of tasks, namely, “working.”

We next split the participants into two age groups according to the median age (36 years). We found that for the younger participants (age<36, N=25), all sound types were superior to silence for producing elevated focus levels (Fig. 6, supp. Table 1, M=0.14,0.13,0.12, SE=0.04,0.03,0.03, t(24)=3.79,4.49,3.67, p=0.004,0.001,0.005 for Endel, Apple and Spotify respectively) while for the older participants (age>36, N=26), there was no difference between sound and silence. The focus level differences were therefore found to also be age-dependent.

### 3.3 Time series analysis of the focus dynamics reveal differences between all sound types and silence

Exploiting the high temporal resolution of the focus measurements, we compared the focus dynamics to each sound stream that played during the 30 minutes of the Preferred Task (Fig. 7, table 2). When comparing Endel’s soundscapes vs. Silence (Fig. 7A), we found that the focus level elicited by Endel’s soundscape was higher 87% of the time, a separation whose significance started after 2.5 minutes of listening. In addition, although on average there wasn’t a significant difference, the focus level elicited by Apple’s playlist was higher than Silence 60% of the time, starting at 12.5 minutes (Fig. 7C), and the focus level elicited by Spotify’s playlist was higher than Silence 27% of the time, starting at 17 minutes (Fig. 7B). Focus elicited by Endel’s soundscape was higher than Spotify’s playlist in 37% of the time, starting at 6 minutes (Fig. 7D).

**Figure 7.**
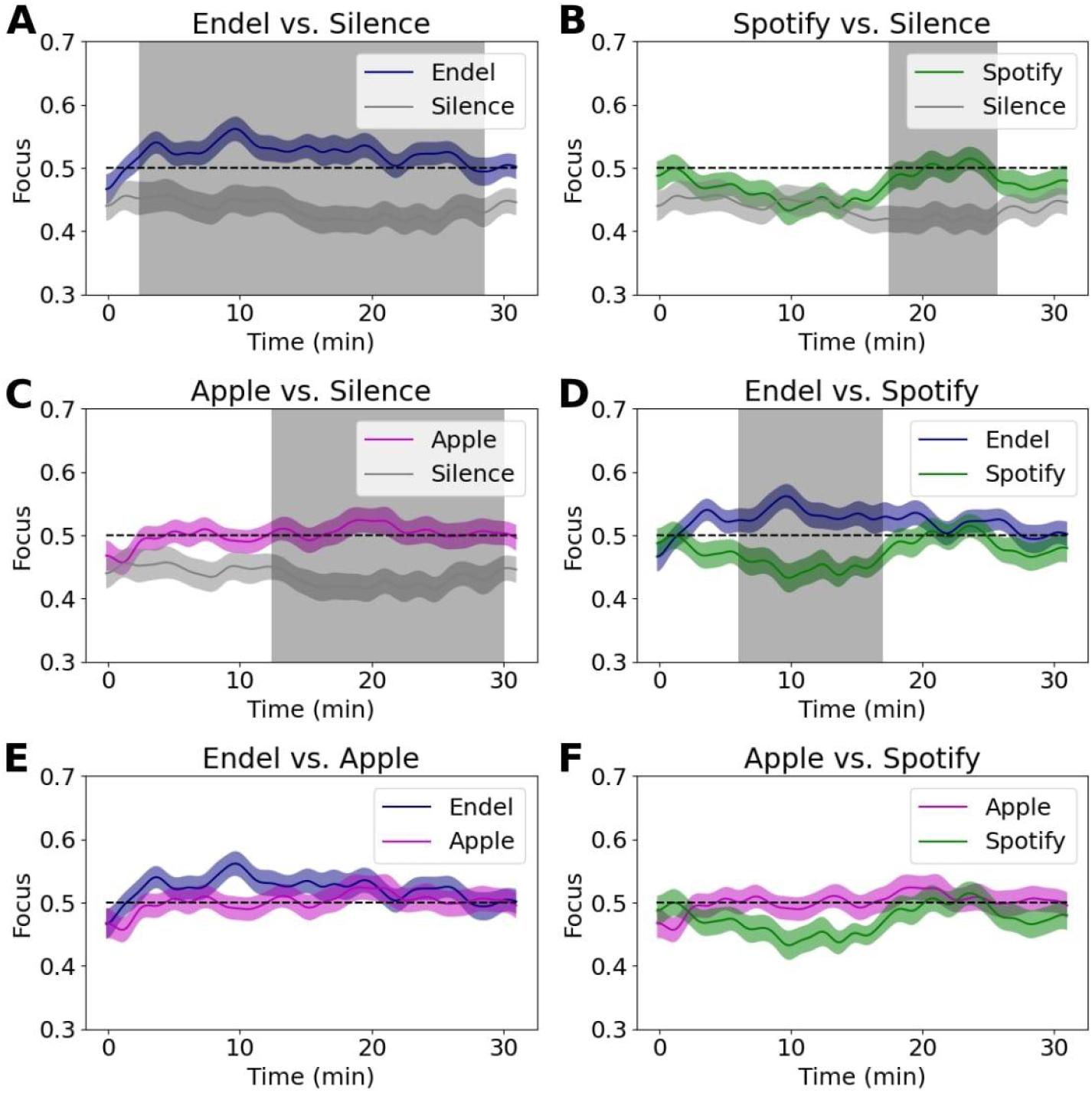
Comparing brain decoded focus dynamics during the 30 minutes of the Preferred Task. Each subfigure shows a comparison between two sound streams, while the gray areas are the timings with a significant difference (p<0.05 corrected, see statistical methods for details).

**Table 2.** Summary of focus time dynamics comparison, showing for each pair the percentage of time and time segments with significant difference (where 100% = 30 minutes).

| Pair | Significant difference<br>(% session) | Significant segments<br>(minutes) |
| --- | --- | --- |
| Endel-Silence | 87% | 2.5-28 |
| Apple-Silence | 60% | 12.5-30 |
| Spotify-Silence | 27% | 17.5-25.5 |
| Endel-Apple | 0% |  |
| Endel-Spotify | 37% | 6-17 |
| Spotify-Apple | 0% |  |

### 3.4 Focus levels in response to sounds can be predicted by the sound’s properties

Seeing that background sound had an effect on focus levels, we go further and ask whether music and soundscapes can be composed according to a formula to increase focus levels. Meaning, can we understand which sound properties drive focus well enough to predict focus levels from only an analysis of the sound properties themselves?

Leveraging the high temporal resolution of the noninvasive brain measurements, we generated a prediction model which predicts the brain-based focus level from sound features extracted from the audio signal. Raw audio files containing the Apple and Spotify sessions were used to extract different sound properties with a running sliding window of 30 seconds. The personalized soundscape session (Endel) was not used in this analysis since the real time streaming did not allow saving the raw audio files that were consistent across participants. Each sound property was treated as a unique feature and checked for its contributory power to the measured average focus level. Supp. Fig. 2 shows the resulting correlations between each sound feature and the brain based focus level. In total, only 20 features of the 136 features evaluated were found to have significant correlations (p<0.05, see Statistical methods).

We next combined multiple sound features to generate a sound data based model that predicts focus levels (see Methods). Figure 8 shows the dynamics of the focus predicted from the audio signal alone which included only properties of the sounds, together with the brain decoded focus that was derived from high resolution electrical brain measurements (Corr=0.7, p<1e-4). Figure 8D shows that if we threshold our dynamics to output a binary prediction (low/high focus), the audio model reaches 88% accuracy in predicting the brain based focus (AUC=0.93).

**Figure 8.**
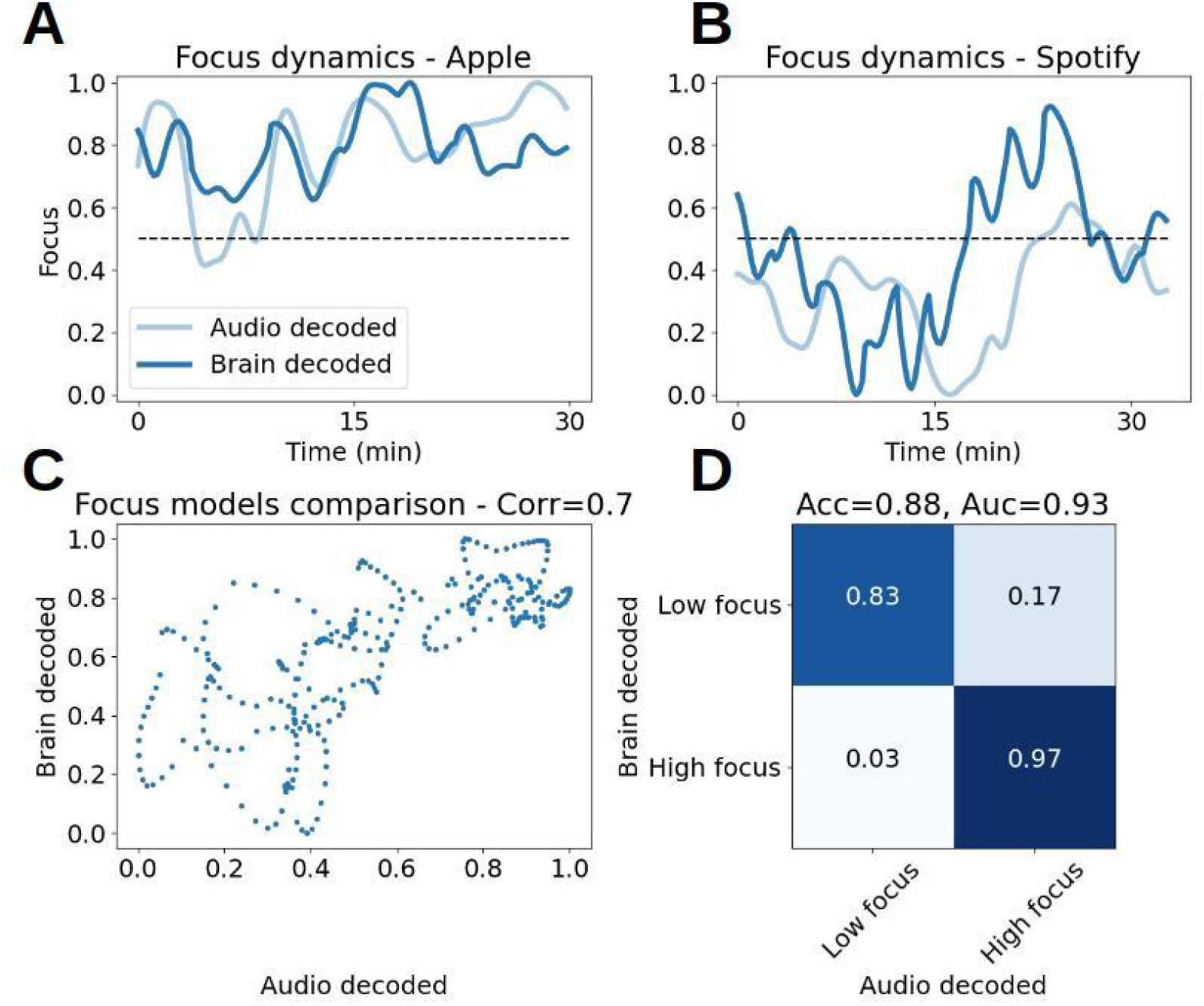
Results of predicting brain decoded focus from audio features. **(A+B)** Dynamics of brain decoded focus (dark blue) and audio decoded focus (light blue), during 30 minutes of the Preferred Task for Apple (A) and Spotify (B). **(C)** Brain decoded focus (y-axis) vs. Audio decoded focus (x-axis) for both playlists (Apple + Spotify). **(D)** Confusion matrix after thresholding the focus predictions to classify between low and high focus. Classification accuracy obtained: 88% (Area under ROC curve: 0.93).

Beyond composing soundscapes for focus, we can also use these prediction models to rate the focus level of a song and assemble successful playlists based on existing songs. To demonstrate this, we compared the song average of the audio decoded output to the brain decoded output. As shown in Figure 9B, there is a correlation of 0.74 between the focus models at the song level (df=16, p=0.0004). Figure 9A shows these averages sorted by the brain-based model.

**Figure 9.**
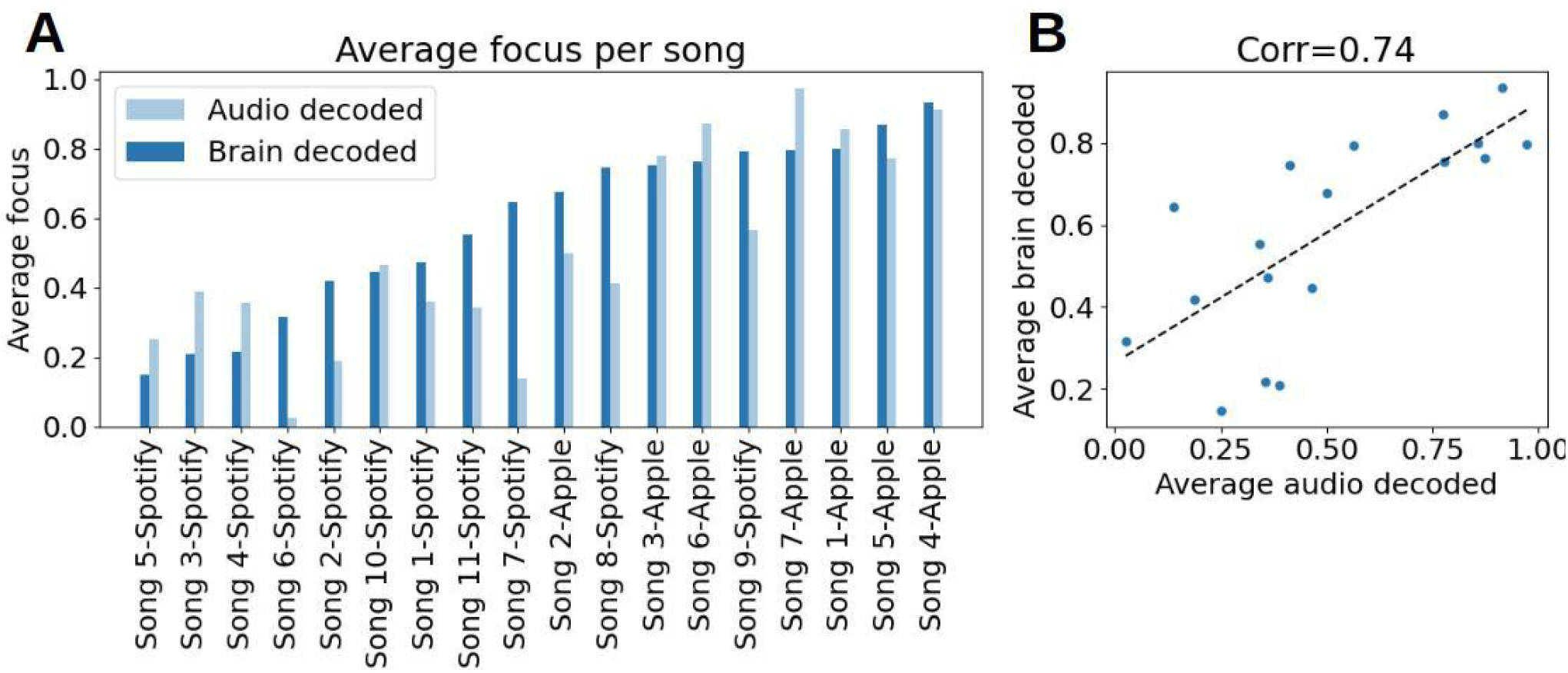
Averaging focus scores for each song. **(A)** Sorted focus scores per song obtained by the brain model (brain decoded - blue), next to the focus obtained by the audio model (audio decoded- light blue). **(B)** Focus scores per song - brain decoded (y axis) vs. audio decoded (x axis). Pearson correlation between them: Corr(16)=0.74, p=0.0004.

### 3.5 Analysis of a sound’s properties can be used to predict its effect on focus

To gain additional insights about the effects of different sound types on human focus, we used the trained audio model, and projected songs and sounds which were not played during the brain recording experiment. Meaning, we obtained their focus score and dynamics based solely on the properties of the sounds they contained. We selected sounds that challenged the validity of the audio model based on their categorical exclusion from the brain recording experiment. A future approach can include these different genres as controls for further brain measurement validation studies. For example, Endel’s soundscapes which are not personalized (taken from the playlist: “[Focus: Calm Clear Morning](https://open.spotify.com/album/473AYNWq9Lxzs26qjnlVDy?si=iRcJWxMSTvGgba8Hd8TQuw&utm_source=copy-link&dl_branch=1&nd=1)”), natural sounds which are commonly used for increasing focus (such as white noise, waves, rain, taken from: https://mc2method.org/white-noise/), and popular songs from other music genres (classical music, electronic, pop, rock, jazz and hip hop) were used.

Figure 10A shows the predicted focus score based on the audio model which took into account only the properties of the sounds themselves. Songs are sorted from the highest focus evoking song (Endel - Three No Paradoxes) to the lowest (Dr. Dre - What’s The Difference). The top two songs are Endel soundscapes which are not personalized, a finding which strengthens our main result since it implies that the high focus scores elicited by Endel’s soundscape was not a byproduct of personalization. Figure 10B shows the sorted focus scores averaged across genres, where notably sounds from classical music and natural sounds contained properties that predicted high focus levels. In contrast, pop and hip-hop songs predicted relatively low focus scores. Although we do not have ground truth focus labels for these songs based on real human brain data, given the relatively high scores of the sounds which are known to have generated increased focus in the experimental data, we can conclude that there is a consistent validity to the model. Future research will gather ground truth labels for these songs and evaluate the model mathematically in this context.

**Figure 10.**
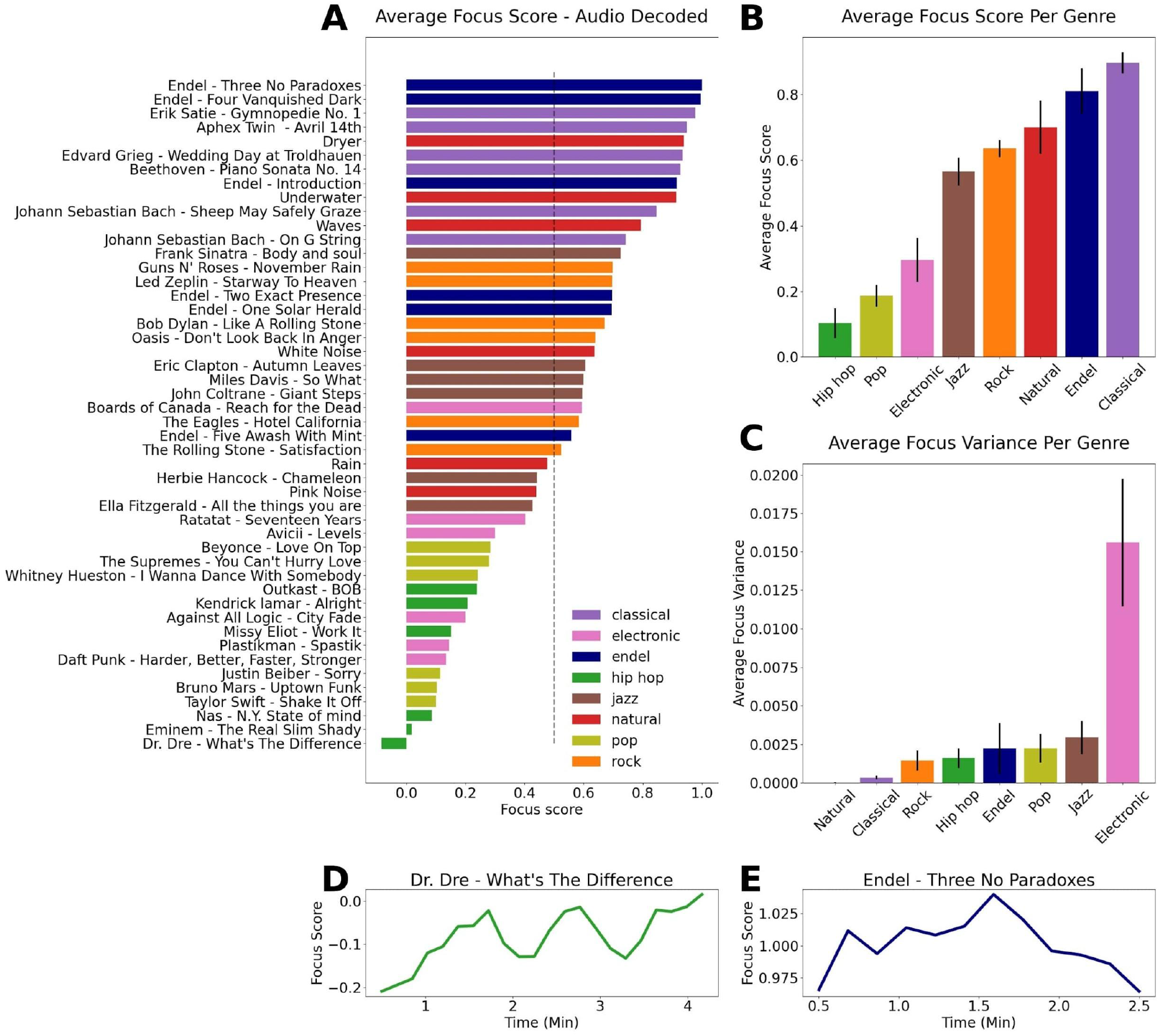
Projecting new songs into the trained audio model. **(A)** Sorted focus scores per song obtained by the audio model, colored by genre. **(B)** Average focus score per genre, sorted from the genre with the lowest score (Hip-hop) to the highest (Classical). **(C)** Average focus variance per genre, sorted from the genre with the lowest variance (Natural) to highest (Electronic). **(D-E)** Focus dynamics for the song with the lowest focus score (D) and the highest (E).

Analyzing the average within-song variance across different genres reveals that the model predicts the largest variance on average for electronic sounds (Fig. 10C), while the lowest variance was found for natural sounds. The variance can be interpreted as a range of focus dynamics, and indeed the focus dynamics of the electronic sounds might change dramatically during a song (supp. Fig3), confirming the need for a tool which outputs dynamics with a high temporal resolution when studying such sound content. Figure 10D-E shows the focus dynamics for the song with the lowest focus evoking score and the highest. The dynamics for all songs can be seen in supp. Fig.3.

## 4 Discussion

“The soundscape of the world is changing. Modern man is beginning to inhabit a world with an acoustical environment radically different from any he has hitherto known” said the composer R. Murray Schafer, presaging the time we live in now when the sounds available to us continue to multiply by the day.

As we have an increasing number of options to modulate our auditory lives by, a handful of take-aways from this study standout:

### 4.1 Brain-based measurement of focus is possible “in-the-wild”

Although the effects of sound and music on the human brain can be subtle in measured brain signals when judging by the changes produced in raw electromagnetic currents, they are robust and highly quantifiable with effectively-trained algorithms, as shown here. Classifying emotional and attentional responses is particularly useful when done at the sub-second temporal resolution which allows one to track dynamics continuously over time at the same timescale as the brain functions that impact perception and behavior. In this study we demonstrated that noninvasive brain decoding technology is able to deliver this needed resolution with a high degree of accuracy (approximately 80% match to self-report, Fig 5C.). Since there are inherent biases in the subjective self-report for experience (Kahneman et al., 1999; Mauss & Robinson, 2009), when mapping physiological signals to self-reported experiences, as done here, there is an upper boundary for accuracy beyond which the model will over fit to the self-reported values and incorrectly represent the information observed in physiological signals. According to a recent review (Larradet et al., 2020) which summarizes multiple peer-reviewed studies that predict self-reported emotions from physiological signals, the average accuracy reported was ~82%. Given this average and the experimental conditions here (small amount of sensors, at home recordings, simple self-report scales), the achieved accuracy was satisfactory for drawing deeper conclusions on sound properties, and aligned with state-of-the-art emotion recognition accuracies in the context of sound as a stimulus (Tripathi et al., 2017, 81.41% and 73.35% for 2 classes of Valence and Arousal respectively).

A key benefit of the current approach is that this method of high temporal resolution brain measurement can be performed reliably outside of traditional laboratories. In this current study not a single laboratory or facility was used for data acquisition. Instead, 18-65 year olds across the U.S. received a technology kit in the mail and experienced music playlists and personalized soundscapes while they recorded their own brain signals from the comfort of their own homes at times of day of their choosing. In other words, in their natural habitat, at their own pace, which lends the research a rare ecological validity.

### 4.2 Focus is increased most by personalized, engineered soundscapes

Within the at-home environment of this study, personalized, engineered soundscapes were found to be the best at increasing participant’s focus levels (Fig. 6A). After 2.5 minutes, on average, listeners of the personalized soundscapes evaluated experienced a meaningful increase in their focus level, while for music playlists it took approximately 15 minutes to gain a similarly appreciable increase (Fig. 7). The audio effect on focus levels was found to be task dependent, where soundscapes increased focus levels in participants who were working (Fig 6B). For participants who were not working, no significant difference was found. This result suggests that willful orientation of attention towards work tasks may have created a brain context especially suited to modification by sound. While engaged in work, participants may also have been more prone to distraction and thus more impacted by the positive uplift of sound compared to reading or playing a game which may have contained more intrinsic motivation to stay focused on.

One limitation of this current study is that it did not allow us to disentangle the effects of personalization of sounds on the listener, since pre-recorded soundscapes were not tested. Equivalently, a comparison of personalized soundscapes to personalized music playlists, where audiences either made their own playlist for focus or were allowed to skip songs whenever they wanted, will likely contribute to a fuller understanding of how sound properties correlate with emotion and attention changes. Follow-up research will incorporate these variables. An additional limitation was the inability to reach conclusions regarding gender-dependent effects which was at least partially due to this study’s imbalanced data set. Despite efforts to recruit a balanced group of participants, enrollment was done on a rolling basis and in the end the female subgroup was statistically underpowered. In future research, especially for closed loop, real-time testing, balanced participant sets will be important for reaching more detailed conclusions.

### 4.3 Sound preferences and focus effects vary between people

It is important to emphasize that the results reported here are sound effects on the average focus levels across a population, and that there was a large variance in this effect between participants. Evidence for this large variance can be seen in Supp Fig. 3 and in the age dependency effect (Fig 6D-E), where for the younger audience, all sounds increased focus while for the older audience, the sounds did not have any effect. These results are consistent with other studies showing personal preferences are critical for the improvements possible by sounds (Cassidy & Macdonald, 2009; Huang & Shih, 2011; Mori et al., 2014). Due to this variety observed together with the highest focus being elicited by the personalized soundscapes, a next step will include closed-loop selections of sounds, where iterative sound testing is used per person to identify the significant parameters for maximizing focus for that person.

Personalized soundscapes specifically, and personalized audio in general, should be investigated further for their capacity to increase productivity, creativity and well-being as these attributes of human experience are associated with one’s ability to focus. For clinical populations as well, for example children with ADHD, the tailoring of sounds for this purpose of increased focus can be particularly impactful. It is possible that the seamlessness of the personalized soundscapes tested here, which played continuously without gaps in the sound like the music playlists had between songs, was a critical part of the observed effect on focus. At every juncture of the experience there is more to be learned, but at a high level, a main lesson of this study is that there is a strong need for personalization of sound in order to most effectively achieve functional goals like increasing focus.

### 4.4 Brain decoded focus data enabled a new predictive model based on sound data alone

Leveraging the high temporal resolution of the brain decoded dynamics, a focus prediction model based on the physical properties of sounds was successfully trained, resulting in an accuracy score of 88% in predicting the brain decoded focus score from an audio decomposition that assessed 136 different properties of sounds as unique features (Fig. 8). This model enabled a further examination of how sounds and different genres effects focus and allowed testing additional conditions, such as pre-recorded soundscapes and commonly used background sounds (e.g. white noise), as well as other genres (pop, rock, jazz, etc). We found that the model predicted the highest focus score to classical music, followed by engineered soundscapes and natural sounds. These results complement previous studies which showed natural sounds and classical music are beneficial for learning and concentration (Angwin et al., 2017; Davies, 2000; DeLoach et al., 2015; H. Liu et al., 2021).

In contrast, the models predicted that genres such as pop and hip-hop produce lower focus scores (Fig 10A-B). It is possible that these sounds contain more distractors that attract attention away from other objects of attention, or that they contain types of sounds that the brain requires more resources to process (depending on familiar patterns, surprises and more), leading to less resources available to perform other tasks. Sounds in these genres may also activate the reward system more (Gold et al., 2019; Salimpoor et al., 2015), which can increase motivation and improve learning of the songs themselves rather than orientation towards other tasks. These sounds may be optimal for driving focus on the songs in other words, rather than focus on other things. Understanding the brain mechanisms that underlie the modified focus is beyond the scope of this work, but the mapping found here can potentially provide fruitful directions for future brain imaging experiments that are equipped to answer these questions.

The analysis here demonstrates a process in which we utilize the temporal resolution of the brain sensing technology to generate a product where the neurotechnology is eventually out of the loop, resulting in a stand alone sound model which gets as an input a raw audio file and outputs a predicted focus score. This model can be used independently to generate focus playlists or to compose optimal soundscapes, and can further be improved by expanding to populations outside the U.S. and different age groups.

Pythagoras, who first identified the mathematical connection between a string’s length and it’s pitch, believed that the whole cosmos was a form of musical composition (James, 1995). We too see the rich mathematical models obtained in this study, by mapping sound properties to human experience, as a glimpse into the natural laws governing how we feel and think. The better these laws can be understood, the more empowered individuals will be to modulate their sound environments to suit their goals and states of mind. There remains much to figure out. While we as a species continue to cause a “shift in the sensorium,” we simultaneously experience that shift all over daily life and it is not clear where we as a species are headed. This study showed that sounds have a distinct effect on our focus, and paves the way for designing sounds to help us focus better in the future.

## 5 Conclusions

We studied the effects of sound on human focus levels using noninvasive brain decoding technology and to gain a better understanding of the optimal sound properties for increasing focus levels in listeners. We combined a custom app, portable EEG-measuring headbands, and brain decoding technology that enabled us to obtain high temporal resolution focus dynamics from participants at home. Using the brain decoded focus dynamics, we then analyzed how various properties of sound affected focus levels in different tasks.

We found that while performing a self-paced task for a long period of time (such as working), personalized soundscapes increased focus the most relative to silence. Curated playlists of pre-recorded songs by Apple and Spotify also increased focus during specific time intervals, especially for the youngest audience demographic. Large variance in response profiles across participants, together with task and age dependent effects, suggest that personalizing sounds in real-time may be the best strategy overall for producing focus in a given listener.

Finally, we generated a sound property based focus model which successfully predicts the brain decoded focus scores using only an audio file as input. Using this model, we extracted predicted focus scores from new songs based on audio decomposition and performed a genre analysis to develop new intuitions about the findings and the source of focus-producing sound content. We found that based on our model, engineered soundscapes and classical music are the best for increasing focus, while pop and hip-hop music are the worst.

The approach taken here can be adapted to include other emotions (e.g. enjoyment, stress, happiness, etc.), attentional parameters (‘Flow state,’ memory formation, etc.) and can be used to assess additional content as well (e.g. visual, ambient, olfactory, etc.), including interactive gaming and e-learning where personalization and high temporal resolution experience measures may be especially beneficial.

## 6 Data Availability Statement

The dataset for this study is available through an open Git repository ([link](https://github.com/arctop/Modeling-the-effect-of-background-sounds-on-human-focus-using-brain-decoding-technology)). Data includes the brain decoded focus dynamics for each participant together with scripts that run the statistical tests.

## 7 Author Contributions

A.H., R.K, N.E, E.K, and D.F designed the experiment, A.H analyzed the data, R.K, N.E, and D.F. advised on data analysis and statistics. S.K and E.K developed the app and software platform for data collection. A.H and D.F wrote the paper, R.K and E.K revised the paper. All authors approved the work for publication.

## 9 Acknowledgments

We would like to thank Hillel Pratt, Kevin Liu, Alexander Kopanev, Warner Music, Sony Music, Endel, and Universal Music for providing audio content, data, and support in conducting this study and advancing theoretical and applied aspects of the research.

## References

Abiri, R., Borhani, S., Jiang, Y., & Zhao, X. (2019). Decoding Attentional State to Faces and Scenes Using EEG Brainwaves. Complexity, 2019, e6862031. https://doi.org/10.1155/2019/6862031

Angwin, A. J., Wilson, W. J., Arnott, W. L., Signorini, A., Barry, R. J., & Copland, D. A. (2017). White noise enhances new-word learning in healthy adults. Scientific Reports, 7(1), 13045. https://doi.org/10.1038/s41598-017-13383-3

Asif, A., Majid, M., & Anwar, S. M. (2019). Human stress classification using EEG signals in response to music tracks. Computers in Biology and Medicine, 107, 182–196.

Bhatti, A. M., Majid, M., Anwar, S. M., & Khan, B. (2016). Human emotion recognition and analysis in response to audio music using brain signals. Computers in Human Behavior, 65, 267–275. https://doi.org/10.1016/j.chb.2016.08.029

Bird, J. J., Ekart, A., Buckingham, C. D., & Faria, D. R. (2019). Mental emotional sentiment classification with an eeg-based brain-machine interface. Proceedings of TheInternational Conference on Digital Image and Signal Processing (DISP’19).

Broday-Dvir, R., Grossman, S., Furman-Haran, E., & Malach, R. (2018). Quenching of spontaneous fluctuations by attention in human visual cortex. NeuroImage, 171, 84–98. https://doi.org/10.1016/j.neuroimage.2017.12.089

Brotzer, J. M., Mosqueda, E. R., & Gorro, K. (2019). Predicting emotion in music through audio pattern analysis. IOP Conference Series: Materials Science and Engineering, 482, 012021. https://doi.org/10.1088/1757-899X/482/1/012021

Cassidy, G., & Macdonald, R. (2009). The effects of music choice on task performance: A study of the impact of self-selected and experimenter-selected music on driving game performance and experience. Musicae Scientiae, 13(2), 357–386. https://doi.org/10.1177/102986490901300207

Chanda, M. L., & Levitin, D. J. (2013). The neurochemistry of music. Trends in Cognitive Sciences, 17(4), 179–193. https://doi.org/10.1016/j.tics.2013.02.007

Cheung, V. K. M., Harrison, P. M. C., Meyer, L., Pearce, M. T., Haynes, J.-D., & Koelsch, S. (2019). Uncertainty and Surprise Jointly Predict Musical Pleasure and Amygdala, Hippocampus, and Auditory Cortex Activity. Current Biology, 29(23), 4084–4092.e4. https://doi.org/10.1016/j.cub.2019.09.067

Chou, P. T.-M. (2010). Attention Drainage Effect: How Background Music Effects Concentration in Taiwanese College Students. Journal of the Scholarship of Teaching and Learning, 10(1), 36–46.

Cowen, A. S., & Keltner, D. (2017). Self-report captures 27 distinct categories of emotion bridged by continuous gradients. Proceedings of the National Academy of Sciences, 114(38), E7900–E7909.

Cunningham, S., Ridley, H., Weinel, J., & Picking, R. (2020). Supervised machine learning for audio emotion recognition. Personal and Ubiquitous Computing, 1–14.

Davies, M. A. (2000). Learning … the Beat Goes on. Childhood Education, 76(3), 148–153. https://doi.org/10.1080/00094056.2000.10522096

Davis, W. B., & Thaut, M. H. (1989). The Influence of Preferred Relaxing Music on Measures of State Anxiety, Relaxation, and Physiological Responses 1. Journal of Music Therapy, 26(4), 168–187. https://doi.org/10.1093/jmt/26.4.168

de la Mora Velasco, E., & Hirumi, A. (2020). The effects of background music on learning: A systematic review of literature to guide future research and practice. Educational Technology Research and Development, 68(6), 2817–2837. https://doi.org/10.1007/s11423-020-09783-4

DeLoach, A. G., Carter, J. P., & Braasch, J. (2015). Tuning the cognitive environment: Sound masking with “natural” sounds in open-plan offices. The Journal of the Acoustical Society of America, 137(4), 2291–2291. https://doi.org/10.1121/1.4920363

Faller, J., Cummings, J., Saproo, S., & Sajda, P. (2019). Regulation of arousal via online neurofeedback improves human performance in a demanding sensory-motor task. Proceedings of the National Academy of Sciences, 116(13), 6482–6490.

Gao, C., Fillmore, P., & Scullin, M. K. (2020). Classical music, educational learning, and slow wave sleep: A targeted memory reactivation experiment. Neurobiology of Learning and Memory, 171, 107206. https://doi.org/10.1016/j.nlm.2020.107206

Giannakopoulos, T. (2015). pyAudioAnalysis: An Open-Source Python Library for Audio Signal Analysis. PLOS ONE, 10(12), e0144610. https://doi.org/10.1371/journal.pone.0144610

Gold, B. P., Mas-Herrero, E., Zeighami, Y., Benovoy, M., Dagher, A., & Zatorre, R. J. (2019). Musical reward prediction errors engage the nucleus accumbens and motivate learning. Proceedings of the National Academy of Sciences, 116(8), 3310–3315. https://doi.org/10.1073/pnas.1809855116

González, V. M., Robbes, R., Góngora, G., & Medina, S. (2015). Measuring Concentration While Programming with Low-Cost BCI Devices: Differences Between Debugging and Creativity Tasks. In D. D. Schmorrow & C. M. Fidopiastis (Eds.), Foundations of Augmented Cognition (pp. 605–615). Springer International Publishing. https://doi.org/10.1007/978-3-319-20816-9_58

Hallam, S., Price, J., & Katsarou, G. (2002). The Effects of Background Music on Primary School Pupils’ Task Performance. Educational Studies, 28(2), 111–122. https://doi.org/10.1080/03055690220124551

Hamadicharef, B., Zhang, H., Guan, C., Wang, C., Phua, K. S., Tee, K. P., & Ang, K. K. (2009). Learning EEG-based spectral-spatial patterns for attention level measurement. 2009 IEEE International Symposium on Circuits and Systems, 1465–1468.

Hizlisoy, S., Yildirim, S., & Tufekci, Z. (2021). Music emotion recognition using convolutional long short term memory deep neural networks. Engineering Science and Technology, an International Journal, 24(3), 760–767. https://doi.org/10.1016/j.jestch.2020.10.009

Hu, J. (2017). Automated Detection of Driver Fatigue Based on AdaBoost Classifier with EEG Signals. Frontiers in Computational Neuroscience, 0. https://doi.org/10.3389/fncom.2017.00072

Huang, R.-H., & Shih, Y.-N. (2011). Effects of background music on concentration of workers. Work, 38(4), 383–387. https://doi.org/10.3233/WOR-2011-1141

Huron, D. B. (2006). Sweet Anticipation: Music and the Psychology of Expectation. MIT Press.

James, J. (1995). The Music of the Spheres: Music, Science, and the Natural Order of the Universe. Copernicus. https://www.springer.com/gp/book/9780387944746

Jung, T.-P., Makeig, S., Stensmo, M., & Sejnowski, T. J. (1997). Estimating alertness from the EEG power spectrum. IEEE Transactions on Biomedical Engineering, 44(1), 60–69.

Kahneman, D., Diener, E., & Schwarz, N. (1999). Well-being: Foundations of hedonic psychology. Russell Sage Foundation.

Kumar, S., von Kriegstein, K., Friston, K., & Griffiths, T. D. (2012). Features versus feelings: Dissociable representations of the acoustic features and valence of aversive sounds. Journal of Neuroscience, 32(41), 14184–14192.

Larradet, F., Niewiadomski, R., Barresi, G., Caldwell, D. G., & Mattos, L. S. (2020). Toward Emotion Recognition From Physiological Signals in the Wild: Approaching the Methodological Issues in Real-Life Data Collection. Frontiers in Psychology, 11, 1111. https://doi.org/10.3389/fpsyg.2020.01111

Larsen, R. J., & Diener, E. (1992). Promises and problems with the circumplex model of emotion.

Levitin, D. J. (2006). This is your brain on music: The science of a human obsession. Penguin.

Levitin, D. J., Chordia, P., & Menon, V. (2012). Musical rhythm spectra from Bach to Joplin obey a 1/f power law. Proceedings of the National Academy of Sciences, 109(10), 3716–3720. https://doi.org/10.1073/pnas.1113828109

Lin, Y.-P., Jao, P.-K., & Yang, Y.-H. (2017). Improving Cross-Day EEG-Based Emotion Classification Using Robust Principal Component Analysis. Frontiers in Computational Neuroscience, 0. https://doi.org/10.3389/fncom.2017.00064

Liu, H., He, H., & Qin, J. (2021). Does background sounds distort concentration and verbal reasoning performance in open-plan office? Applied Acoustics, 172, 107577. https://doi.org/10.1016/j.apacoust.2020.107577

Liu, N.-H., Chiang, C.-Y., & Chu, H.-C. (2013). Recognizing the Degree of Human Attention Using EEG Signals from Mobile Sensors. Sensors (Basel, Switzerland), 13(8), 10273–10286. https://doi.org/10.3390/s130810273

Mauss, I. B., & Robinson, M. D. (2009). Measures of emotion: A review. Cognition and Emotion, 23(2), 209–237. https://doi.org/10.1080/02699930802204677

Micoulaud-Franchi, J.-A., Geoffroy, P. A., Fond, G., Lopez, R., Bioulac, S., & Philip, P. (2014). EEG neurofeedback treatments in children with ADHD: An updated meta-analysis of randomized controlled trials. Frontiers in Human Neuroscience, 0. https://doi.org/10.3389/fnhum.2014.00906

Mori, F., Naghsh, F. A., & Tezuka, T. (2014). The Effect of Music on the Level of Mental Concentration and its Temporal Change. Proceedings of the 6th International Conference on Computer Supported Education - Volume 1, 34–42. https://doi.org/10.5220/0004791100340042

Nia, H. T., Jain, A. D., Liu, Y., Alam, M.-R., Barnas, R., & Makris, N. C. (2015). The evolution of air resonance power efficiency in the violin and its ancestors. Proceedings of the Royal Society A: Mathematical, Physical and Engineering Sciences, 471(2175), 20140905. https://doi.org/10.1098/rspa.2014.0905

Perez-Valero, E., Vaquero-Blasco, M. A., Lopez-Gordo, M. A., & Morillas, C. (2021). Quantitative Assessment of Stress Through EEG During a Virtual Reality Stress-Relax Session. Frontiers in Computational Neuroscience, 0. https://doi.org/10.3389/fncom.2021.684423

Rebolledo-Mendez, G., Dunwell, I., Martínez-Mirón, E. A., Vargas-Cerdán, M. D., de Freitas, S., Liarokapis, F., & García-Gaona, A. R. (2009). Assessing NeuroSky’s Usability to Detect Attention Levels in an Assessment Exercise. In J. A. Jacko (Ed.), Human-Computer Interaction. New Trends (pp. 149–158). Springer. https://doi.org/10.1007/978-3-642-02574-7_17

Sacks, O. (2010). Musicophilia: Tales of music and the brain. Vintage Canada.

Salimpoor, V. N., Zald, D. H., Zatorre, R. J., Dagher, A., & McIntosh, A. R. (2015). Predictions and the brain: How musical sounds become rewarding. Trends in Cognitive Sciences, 19(2), 86–91. https://doi.org/10.1016/j.tics.2014.12.001

Schreiber, C. A., & Kahneman, D. (2000). Determinants of the remembered utility of aversive sounds. Journal of Experimental Psychology: General, 129(1), 27.

Shahabi, H., & Moghimi, S. (2016). Toward automatic detection of brain responses to emotional music through analysis of EEG effective connectivity. Computers in Human Behavior, 58, 231–239. https://doi.org/10.1016/j.chb.2016.01.005

Shih, Y.-N., Huang, R.-H., & Chiang, H.-Y. (2012). Background music: Effects on attention performance. Work, 42(4), 573–578. https://doi.org/10.3233/WOR-2012-1410

Tripathi, S., Acharya, S., Sharma, R. D., Mittal, S., & Bhattacharya, S. (2017). Using Deep and Convolutional Neural Networks for Accurate Emotion Classification on DEAP Dataset. Twenty-Ninth IAAI Conference.

Tuckute, G., Hansen, S. T., Kjaer, T. W., & Hansen, L. K. (2021). Real-Time Decoding of Attentional States Using Closed-Loop EEG Neurofeedback. Neural Computation, 33(4), 967–1004. https://doi.org/10.1162/neco_a_01363

Vempala, N. N., & Russo, F. A. (2012). Predicting emotion from music audio features using neural networks. Proceedings of the 9th International Symposium on Computer Music Modeling and Retrieval (CMMR), 336–343.

Washburne, C. (2020). “More Cowbell”: Latin Jazz in the Twenty-First Century. In Latin Jazz. Oxford University Press. https://doi.org/10.1093/oso/9780195371628.003.0007

Yang, Y.-H., Lin, Y.-C., Su, Y.-F., & Chen, H. H. (2008). A Regression Approach to Music Emotion Recognition. IEEE Transactions on Audio, Speech, and Language Processing, 16(2), 448–457. https://doi.org/10.1109/TASL.2007.911513

Zald, D. H., & Pardo, J. V. (2002). The neural correlates of aversive auditory stimulation. Neuroimage, 16(3), 746–753.

