## Supplementary for "Modeling The Effect of Background Sounds on Human Focus Using Brain Decoding Technology"

### *Supplementary Material*

#### 1 Supplementary Figures and Tables

##### 1.1 Supplementary Figures

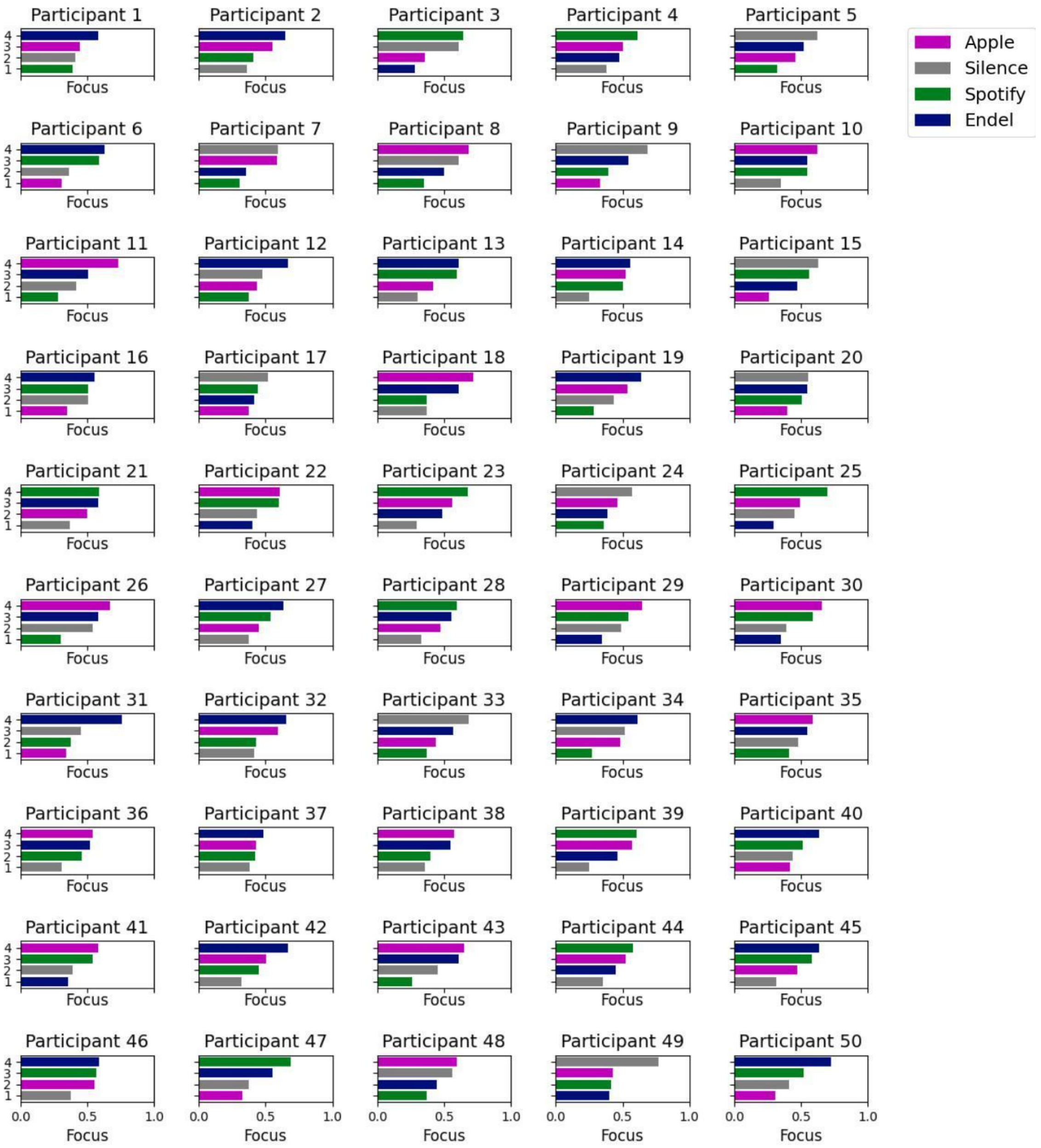


**Supplementary Figure 1. Average Brain decoded focus levels for all participants in all sessions' Preferred Task.** For each participant, the sessions are sorted from their highest average focus level (4) to the lowest (1), with colors representing the different experimental audio streams. The distribution of highest sessions is presented in Fig. 6B.

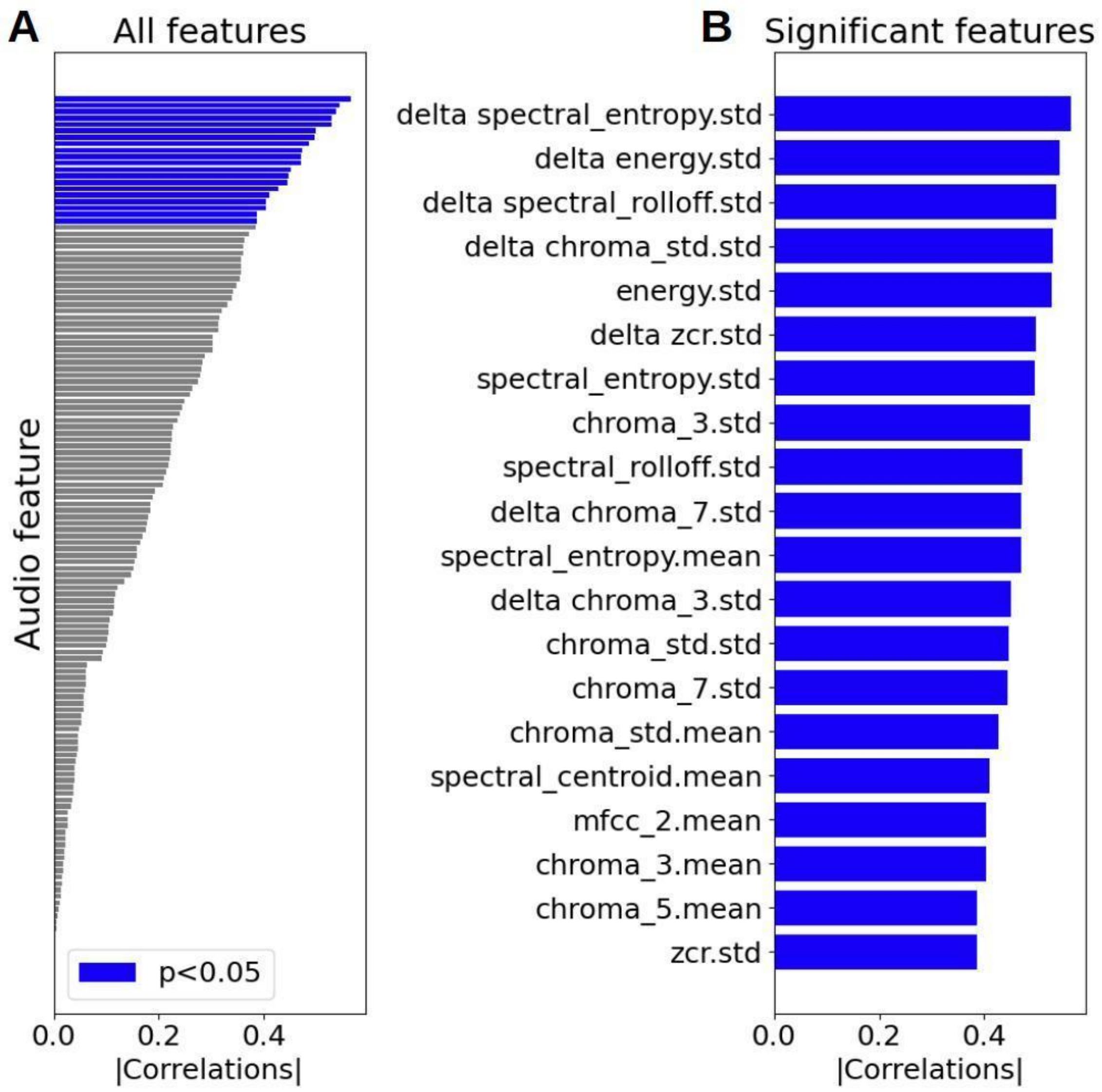


**Supplementary Figure. 2. Single features correlations (absolute value) between audio features extracted from music playlists and listener brain decoded focus**. A. Correlations for all extracted features (see Methods). Significant features are colored in blue. B. Only significant features (|Corr|>0.39) are named at right, see Statistical methods for details).

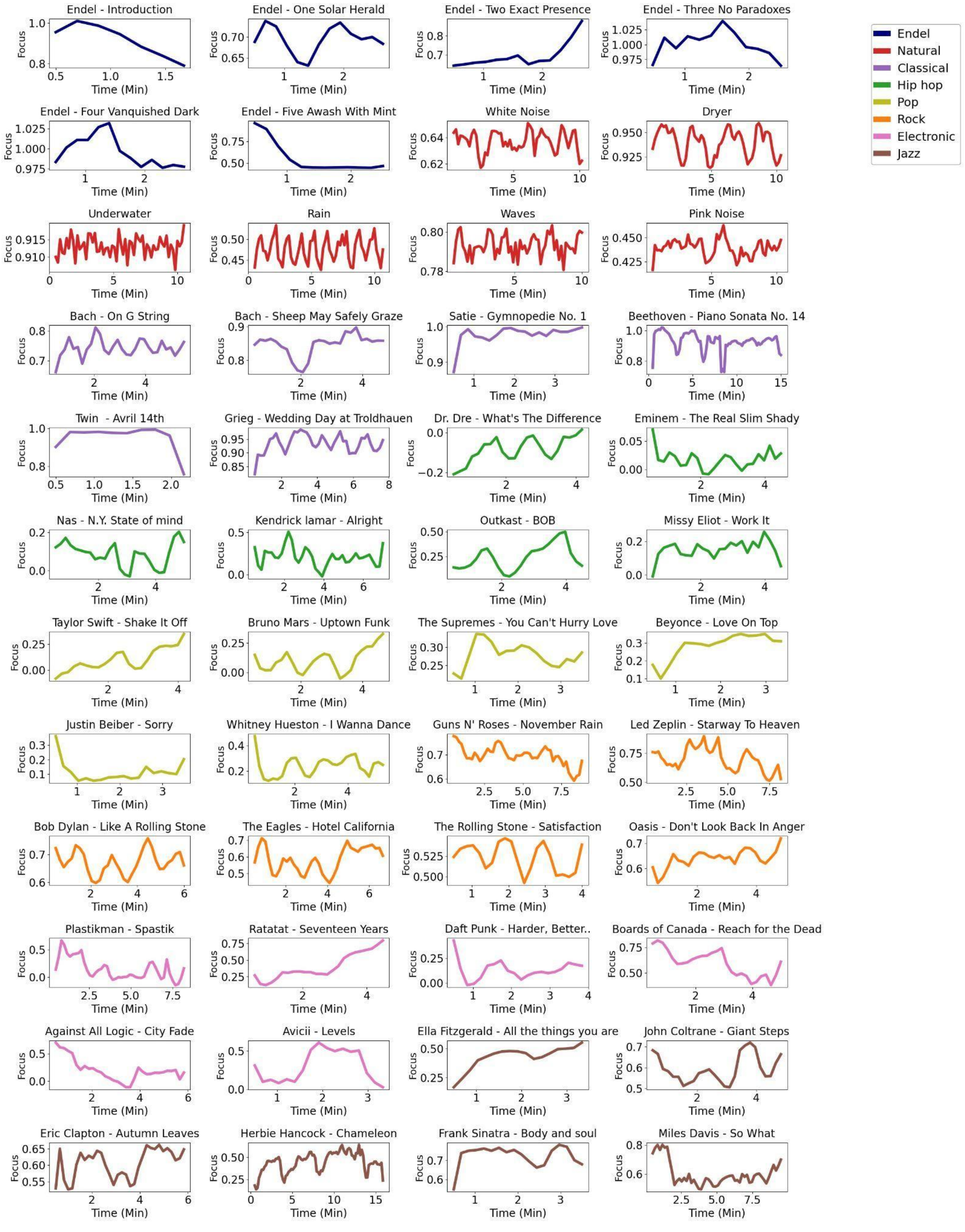


**Supplementary Figure 3. Audio decoded focus dynamics for all songs.** Each color represents a different genre according to the legend on the right. The properties of sound in each audio file were used exclusively to simulate listener focus dynamics - no brain measurements from real audiences were required

#### 1.2 Supplementary Tables

**Supplementary Table 1. Results of post-hoc statistical tests, comparing all pairs of audio streams' focus levels for all and for each subgroup.** P values are corrected using the Holm-Bonferroni method. Significant differences are colored in grey (p(holm)<0.05) and as long as the Anova test (Table 1) revealed significance differences..

| Group | Pair | Avg | Ste | t | df | p (holm) |
|---|---|---|---|---|---|---|
| **All** | **Silence-Endel** | **-0.090** | **0.027** | **-3.38** | **50** | **0.008** |
| | Silence-Apple | -0.063 | 0.026 | -2.37 | 50 | 0.107 |
| | Silence-Spotify | -0.036 | 0.028 | -1.24 | 50 | 0.653 |
| | Endel-Apple | 0.027 | 0.024 | 1.13 | 50 | 0.653 |
| | Endel-Spotify | 0.054 | 0.025 | 2.13 | 50 | 0.153 |
| | Spotify-Apple | -0.027 | 0.026 | 1.03 | 50 | 0.653 |
| **Working** | **Silence-Endel** | **-0.119** | **0.036** | **-3.26** | **25** | **0.017** |
| | Silence-Apple | -0.058 | 0.034 | -1.68 | **25** | 0.317 |
| | Silence-Spotify | -0.072 | 0.036 | -1.172 | **25** | 0.292 |
| | Endel-Apple | 0.061 | 0.033 | 1.155 | **25** | 0.317 |
| | Endel-Spotify | 0.047 | 0.039 | 2.131 | **25** | 0.487 |
| | Spotify-Apple | -0.014 | 0.034 | 1.015 | **25** | 0.687 |
| **Not Working** | **Silence-Endel** | **-0.060** | **0.038** | **-1.552** | **24** | **0.447** |
| | Silence-Apple | -0.067 | 0.041 | -1.650 | **24** | 0.447 |
| | Silence-Spotify | 0.002 | 0.044 | 0.045 | **24** | 1.000 |
| | Endel-Apple | -0.007 | 0.034 | -0.202 | **24** | 1.000 |
| | Endel-Spotify | 0.062 | 0.033 | 1.870 | **24** | 0.442 |
| | Spotify-Apple | 0.069 | 0.037 | 1.838 | **24** | 0.442 |
| **Age > 36** | **Silence-Endel** | **-0.044** | **0.037** | **-1.17** | **25** | **1.000** |
| | Silence-Apple | 0.005 | 0.039 | 0.12 | **25** | 1.000 |
| | Silence-Spotify | 0.043 | 0.042 | 1.01 | **25** | 1.000 |
| | Endel-Apple | 0.048 | 0.034 | 1.42 | **25** | 0.840 |
| | Endel-Spotify | 0.086 | 0.029 | 2.91 | **25** | 0.044 |
| | Spotify-Apple | 0.037 | 0.039 | 0.97 | **25** | 1.000 |
| **Age < 36** | **Silence-Endel** | **-0.139** | **0.036** | **-3.79** | **24** | 0.004 |
| | Silence-Apple | -0.133 | 0.029 | -4.49 | **24** | 0.001 |
| | Silence-Spotify | -0.117 | 0.032 | -3.67 | **24** | 0.005 |
| | Endel-Apple | 0.006 | 0.034 | 0.16 | **24** | 1.000 |

| | | | | | | |
|---|---|---|---|---|---|---|
| | Endel-Spotify | 0.021 | 0.042 | 0.51 | **24** | 1.000 |
| | Spotify-Apple | -0.015 | 0.035 | 0.44 | **24** | 1.000 |
